# Phylogeny and evolution of the SARS-CoV-2 spike gene from December 2022 to February 2023

**DOI:** 10.1101/2023.07.23.549423

**Authors:** Hsiao-Wei Kao

## Abstract

**Background:** By the end of 2022, new variants of SARS-CoV-2, such as BQ.1.1.10, BA.4.6.3, XBB, and CH.1.1, emerged with higher fitness than BA.5.

**Methods:** The file (spikeprot0304), which contains spike protein sequences, isolates collected before March, 4, 2023, was downloaded from Global Initiative on Sharing All Influenza Data (GISAID). A total of 188 different spike protein sequences were chosen, of which their isolates were collected from December 2022 to February 2023. These sequences did not contain undetermined amino acid X, and each spike protein sequence had at least 100 identical isolate sequences in GISAID. Phylogenetic trees were reconstructed using IQ-TREE and MrBayes softwares. A median-join network was reconstructed using PopART software. Selection analyses were conducted using site model of PAML software.

**Results:** The phylogenetic tree of the spike DNA sequences revealed that the majority of variants belonged to three major lineages: BA.2 (BA.1.1.529.2), BA.5 (BA.1.1.529.5), and XBB. The median network showed that these lineages had at least six major diversifying centers. The spike DNA sequences of these diversifying centers had the representative accession IDs (EPI_ISL_) of 16040256 (BN.1.2), 15970311 (BA.5), 16028739 (BA.5.11), 16028774 (BQ.1), 16027638 (BQ.1.1.23), and 16044705 (XBB.1.5). Selection analyses revealed 26 amino-acid sites under positive selection. These sites included L5, V83, W152, G181, N185, V213, H245, Y248, D253, S255, S256, G257, R346, R408, K444, V445, G446, N450, L452, N460, F486, Q613, Q675, T883, P1162, and V1264.

**Conclusion:** The spike proteins of SARS-CoV-2 from December 2022 to February 2023 were characterized by a swarm of variants that were evolved from three major lineages: BA.2 (BA.1.1.529.2), BA.5 (BA.1.1.529.5), and XBB. These lineages had at least six diversifying centers. Selection analysis identified 26 amino acid sites were under positive selection. Continued surveillance and research are necessary to monitor the evolution and potential impact of these variants on public health.

## Background

On May 5, 2023, the World Health Organization (WHO) declared that COVID- 19 is no longer a public health emergency of international concern (PHEIC) due to the decreasing trend in COVID-19 deaths, decline in COVID-19-related hospitalizations and intensive care unit admissions, and the high levels of population immunity to SARS-CoV-2 [1].

The Omicron (B.1.1.529) variant was designated as the fifth variant of concern declared by the WHO on November 26, 2021 [2]. A comparison between the B.1.529 variant and the Wuhan-Hu-1 genome sequences revealed 53 nucleotide substitutions. Within these substitutions, 30 were nonsynonymous substitutions located in the spike gene [3, 4]. Additionally, there were six amino acid deletions at positions 69, 70, 143, 144, 145, and 211. Furthermore, three amino acid insertions (EPE) were observed between positions 214 and 215, relative to the amino acid positions in the Wuhan-Hu- 1 spike protein [3, 4].

The major lineages that contributed to the pandemic from 2019 to 2022 were Omicron BA.1, BA.2, BA.3, BA.4, and BA.5 [5]. Recently, new variants have emerged, including BQ.1.1.10, BA.4.6.3, XBB, and CH.1.1, which had higher fitness than BA.5 [6–8]. This higher fitness includes evasion of neutralization drugs and convalescent plasma, even those targeting BA.5 breakthrough infections. The immune escape mechanism of these new variants is primarily attributed to specific mutations at amino acid sites R346, R356, K444, V445, G446, N450, L452, N460, F486, F490, R493, and S494 within the receptor binding domain of the spike protein. These mutations have been observed in at least five different phylogenetic lineages, which suggests that there has been convergent evolution of the receptor binding domain driven by preexisting SARS-CoV-2 humoral immunity [6–8].

In this study, the evolution of the SARS-CoV-2 spike gene between December 2022 and February 2023 was investigated. To summarize the major lineages of SARS-CoV-2 and their spike gene evolution during this period, a phylogenetic tree and median-joining network were reconstructed. Furthermore, to identify amino acid sites that were potentially under positive selection and associated with adaptive changes in the spike gene, the nonsynonymous versus synonymous substitution ratio (dn/ds ratio = ω) was calculated. This was done using the site model in the codeml module of the PAML software [9].

## Methods

### Data collection and analyses

The file “spikeprot0304” containing spike protein sequences was downloaded from the Global Initiative on Sharing All Influenza Data (GISAID) [10]. To filter the sequences, the following criteria were applied using the Bioedit software [11]: the collection days ranged from December 2022 to February 2023, the sequence lengths ranged from 1259 to 1319 amino acids, and sequences without undetermined amino acid X were included. After filtering, a total of 369,809 spike protein sequences were obtained from the “spikeprot0304” file. To determine the number of identical isolate sequences for different spike protein sequences in the GISAID database, the 369809 spike protein sequences were further filtered using different spike protein sequences as references. Ultimately, 188 different spike protein sequences, referred to as protein haplotypes, were obtained. Each protein haplotype consisted of at least 100 identical isolate sequences within the set of 369809 spike protein sequences. For each protein haplotype, one representative accession ID (GIS_ISL_) was selected.

To obtain the DNA sequences corresponding to the 188 spike protein haplotypes, I downloaded the complete genomes of these haplotypes from GISAID using their accession IDs. The downloaded complete genomes comprised the SARS-CoV-2 DNA sequences. I aligned the 188 complete genomes using MAFFT v.7.450 software [12], using the Wuhan-Hu-1 sequence (GenBank accession number: MN908947.3) as the reference sequence. The resulting alignment contained 189 DNA sequences, including the additional Wuhan-Hu-1 sequence. The spike DNA sequences were cut to a new alignment for phylogenetic and selection analyses.

To align the 189 spike DNA sequences, the DNA sequences were first translated into protein sequences using the Bioedit software. The translated protein sequences were then aligned using MAFFT v.7.450 software. Based on the alignment of the protein sequences, the corresponding DNA sequences were aligned using the Dambe software [13].

### Reconstruction of phylogenetic tree and median join network

I used the jmodeltest software [14] to determine the best evolutionary model for the alignment of the spike DNA sequences. To reconstruct phylogenetic tree, I conducted maximum likelihood (ML) and Bayesian analyses using IQ-TREE software [15] and MrBayes software [16], respectively. In ML analysis, the statistical support for the tree topology was assessed using 1000 bootstrap replicates. In BA analysis, the parameters of the likelihood model were set as nst = 6 and rate = invgamma, as determined by jmodeltest. The analysis was run for 10^7^ generations, with a sample frequency of 1000 and a burn-in of 2500. The consensus tree with posterior probability was constructed based on 7500 trees.

I reconstructed a median-join network based on the 189 spike DNA sequences. The lineages of the spike sequences were assigned according to the Pango-lineage nomenclatures [17] in the GISAID. The median network of the 189 spike DNA haplotypes was constructed with PopART software [18]. To enhance the visualization of different lineages in the phylogenetic tree and median-join network, I used Inkscape and PowerPoint to edit the phylogenetic tree and median-join network. In Inkscape, I assigned different colors to the Pango lineages based on hexadecimal codes, while in PowerPoint, I used the corresponding RGB values to color-code the lineages. These editing steps were performed to facilitate the easy identification and differentiation of the various spike protein lineages in the phylogenetic tree and median-join network.

To calculate the genetic distances between the major lineages of SARS-Cov-2, the 189 spike DNA haplotypes were divided into nine major groups: Wuhan-Hu-1, BA.1.1.529.2 (BA.2), BA.1.1.529.4 (BA.4), B.1.1.529.5 (BA.5), XBB.1, XBC, XBF, XBM, and XBZ. The net average distance (the net number of amino acid differences per sequence) was computed for all sequence pairs between these major groups using MEGA11 software [19]. The net average distance between two groups is given by

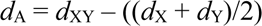

Where, *d*XY is the average distance between groups X and Y, and *d*X and *d*Y are the mean within-group distances [19]. The analysis assumed a uniform rate among sites, and pairwise deletion was used to handle gaps between sequences.

To determine whether specific amino acid sites in the spike proteins of SARS-Cov-2 were under selection, the nonsynonymous versus synonymous substitution ratio (dn/ds ratio = ω) was calculated using the site model in the codeml program of the PAML software [20]. The ω ratio provides information about the balance between nonsynonymous (amino acid-changing) and synonymous (amino acid-preserving) substitutions at each site. A value of ω < 1 suggests purifying (negative) selection, ω = 1 suggests neutral evolution, and ω > 1 suggests positive (diversifying) selection.

Likelihood ratio tests were performed to compare different evolutionary models: M0 (one ratio) versus M3 (discrete), M1a (nearly neutral) versus M2 (selection), and M7 (beta) versus M8 (beta & ω). The Bayes empirical Bayes method was used to calculate posterior probabilities for site classes [21]. If the likelihood ratio test is statistically significant, it suggests that the amino acid sites are under selection. It is important to note that only the 188 spike DNA haplotypes were analyzed in this study. The Wuhan-Hu-1 sequence was not included in the analyses due to the absence of Wuhan-Hu-1 spike protein haplotypes in the GISAID database from December 1, 2012, to February 2013. Amino acid sites with gaps in the spike DNA sequence alignment were deleted because the nonsynonymous versus synonymous substitution value cannot be calculated in the PAML software. The site numbering used the spike protein (protein ID=QHD416.1) of the Wuhan-Hu-1/2019 (GenBank accession number MN908947.3) as the reference for consistency.

## Results

### Characteristics of the spike protein sequences

According to the filtering criteria mentioned, a total of 369809 spike protein sequences were obtained from the spikeprot0304 file. Among these sequences, 221323 isolates were collected in December 2022, 119971 isolates in January 2023, and 28515 isolates in February 2023. No isolate was filtered out in March 2023. The number of isolate sequences versus amino acid lengths of spike protein sequences is as follows: 1710 isolate sequences had 1266 amino acids, 57587 isolate sequences had 1267 amino acids, 216386 isolate sequences had 1268 amino acids, 45463 isolate sequences had 1269 amino acids, 47036 isolate sequences had 1270 amino acids, 547 isolate sequences had 1271 amino acids, 528 isolate sequences had 1272 amino acids, and 253 isolate sequences had 1273 amino acids. Other spike protein sequences with lengths of 1259, 1260, 1261, 1262, 1263, 1264, 1265, 1274, 1275, 1276, 1277, 1281, 1283, or 1319 amino acids had fewer than 72 isolate sequences (Fig. 1). Out of the 189 spike protein haplotypes analyzed, there were 4 haplotypes with 1266 amino acids, 36 haplotypes with 1267 amino acids, 106 haplotypes with 1268 amino acids, 16 haplotypes with 1269 amino acids, 25 haplotypes with 1270 amino acids, one haplotype with 1272 amino acids, and one haplotype with 1273 amino acids. The haplotype with 1273 amino acids is the Wuhan-Hu-1 sequence, but its spike protein haplotype was not found in the GISAID database from December, 2022 to February, 2023.

**Fig 1.**
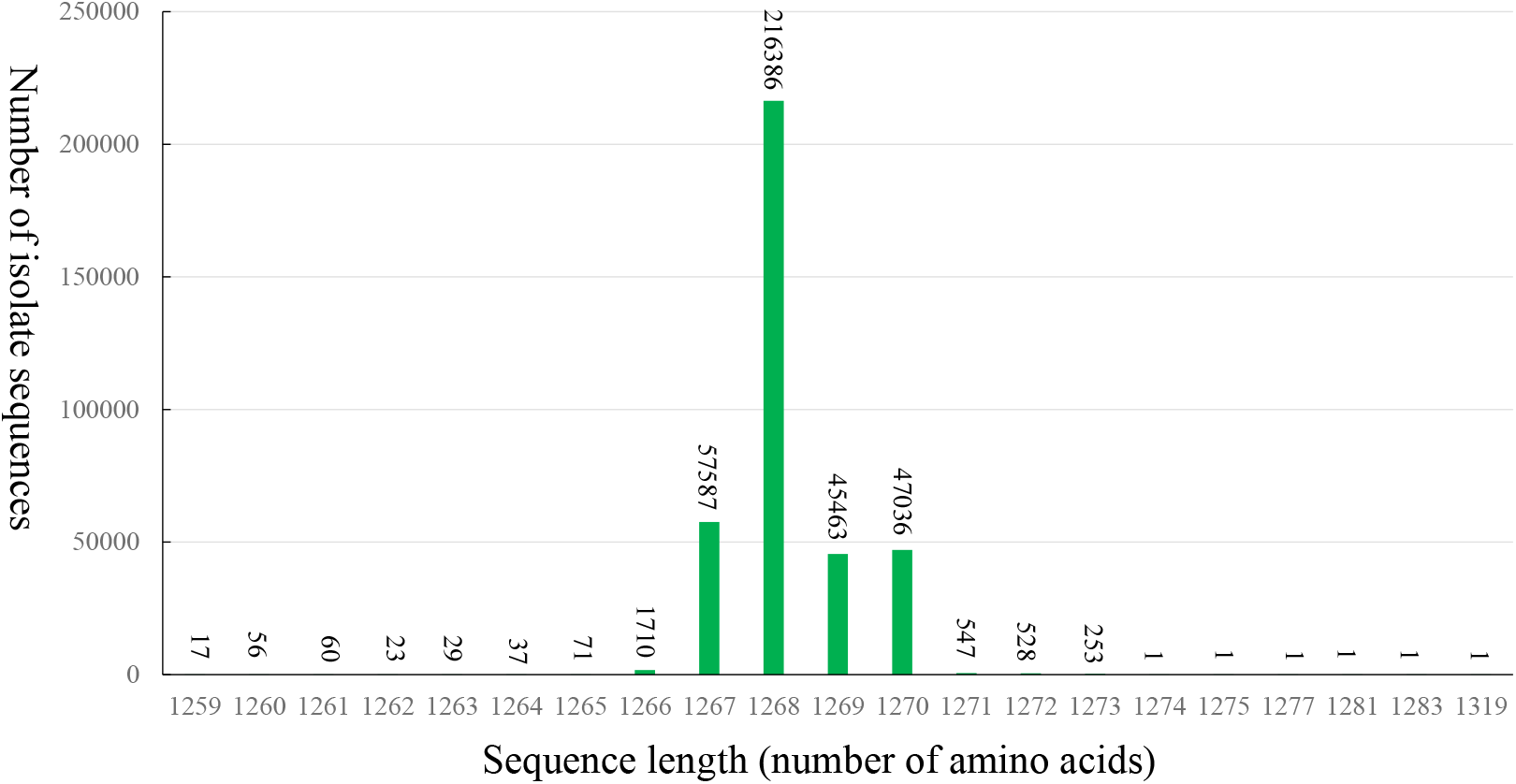
Number of isolate sequences versus different lengths of spike protein in GISAID from December 2022 to February 2023.

### Net average genetic distances of spike proteins between major lineages of SARS- CoV-2

The net average genetic distances of spike protein between Wuhan-Hu-1 and B.1.1.529.2 (BA.2), B.1.1.529.4 (BA.4), B.1.1.529.5 (BA.5), XBB, XBC, XBF, and XBM were 34.54, 31, 37.07, 36.62, 35, 37, 33, and 31 amino acids per sequence, respectively. The net average genetic distances of spike protein between B.1.1.529.2 (BA.2) and B.1.1.529.4 (BA.4), B.1.1.529.5 (BA.5), XBB, XBC, XBF, XBM, and XBZ were 9.41, 7.3, 11.87, 16.71, 1.71, 11.41, and 8.67 amino acids per sequence, respectively. The net average genetic distances of spike protein between B.1.1529.4 (BA.4) and B.1.1.529.5 (BA.5), XBB, XBC, XBF, XBM, and XBZ were 1.99, 13.18, 15, 12, 4, and 4 amino acids per sequence, respectively. The net amino acid differences per sequence of spike protein between B.1.1.529.5 (BA.5) and XBB, XBC XBF, XBM, and XBZ were 11.73, 14.12, 10.52, 3.9, and 1.4 amino acids per sequence, respectively. The net average genetic distances of spike protein between XBB and XBC, XBF, XBM, and XBZ were 19.62, 11.62, 15.18, and 12.93 amino acids per sequence, respectively. The net average genetic distances of spike protein between XBC, and XBF, XBM, and XBZ were 18, 17, 17 amino acids per sequence, respectively. The net average genetic distances of spike protein between XBF and XBM and XBZ was 14 and 12 amino acids per sequence, respectively. The net average genetic distances of spike protein between XBM and XBZ was 6 amino acids per sequence (Table.1).

### Phylogenetic analyses of spike DNA sequences

The phylogenetic tree of 189 spike DNA sequences (Fig. 2) consisted of three major clades. Clade I consisted of lineages or descendants of BQ.1, BF, and DN. It was positioned closer to the root of the tree. Clade II consisted of lineages or descendants of BA.5. It was located between clade I and clade III in the phylogenetic tree. Clade III was further distal to the root compared to clade II and consisted of subclades A, B, C, and D. Subclade A consisted of lineages or descendants of CM. Subclade B encompasses lineages or descendants of CH.1, CA, CV, and BR. Subclade C consisted of lineages or descendants of BN.1. Subclade D consisted of the lineage or descendant of XBB lineages. In the maximum likelihood (ML) analysis, it was found that the sequences BF.1.1 (EPI_ISL_16152392) and BF.7 (EPI_ISL_16080401) within clade I occupied the most basal position when the phylogenetic tree was rooted by the Wuhan-Hu-1 sequence. Statistical analyses, including bootstrap values and posterior probabilities, provided strong support for the monophyly (common ancestry) of clade III and its subclades A, C, and D. A bootstrap value or posterior probability of more than 0.95 indicated a high level of confidence in the grouping of sequences within these clades.

**Fig. 2.**
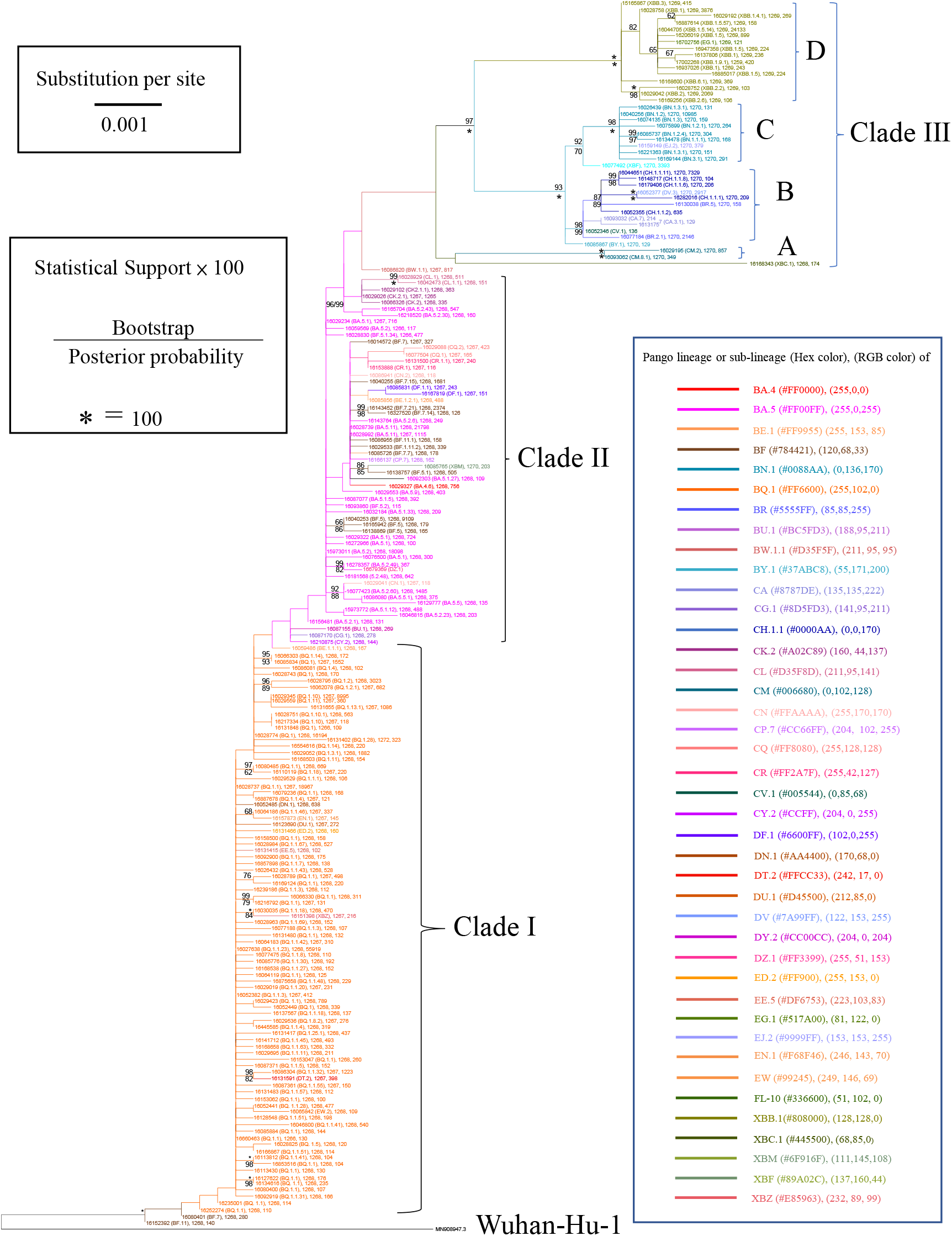
Phylogeny of SARS-CoV-2 spike DNA sequences. The terminal node (leaf) is the GISAID ID of the sequence followed by the lineage name in parentheses, the length of the spike protein, and the number of isolates. Statistical supports are labeled on the branches. The values below 60% are not labeled.

### Median-join network of spike DNA sequences

Median-join network (Fig. 3) showed that the BF.11 (EPI_ISL_16152392) connected to Wuhan-Hu-1 Spike DNA sequences with 29 nucleotide substitutions. The network can be classified into six major clusters, i.e., BQ.1, BA.5, CH.1.1, CM, BN.1 and XBB.1. The BQ.1 cluster had two diversifying centers. In the BQ.1 cluster’s first diversifying center, there were nine haplotypes with the following GISAID accession IDs (EPI_ISL_): 16027638, 16028737, 16029423, 16052382, 16052485, 16064186, 16077475, 16113812, and 16660463. It is worth noting that these nine DNA sequences were considered identical in the analysis because the PopART software only counted nucleotide substitutions and did not count insertions or deletions in the alignment. In the BQ.1 cluster’s second diversifying center, there were seven sequences with GISAID accession IDs (EPI_ISL_) of 16028751, 16028774, 16029345, 16029559, 16052449, 16131848, and 16217334. These sequences also exhibited differences due to insertions and deletions. Among them, the spike protein sequence of EPI_ISL_16028774 was the most abundant, with 16194 isolates recorded in GISAID. The BA.5 cluster consisted of three haplotypes with GISAID accession IDs (EPI_ISL_) of 15973011, 16029234, and 16059569. These three spike DNA sequences also exhibited variations due to insertions and deletions. Among them, the spike protein sequence of EPI_ISL_15973011 was the most abundant, with 18098 isolates recorded in GISAID. The CH.1.1 cluster consisted of six spike DNA haplotypes that had diversified from an unknown haplotype. Among them, the spike protein haplotype (EPI_ISL_16044651) was the most abundant, with 7329 isolates recorded. It differed from the haplotype of EPI_ISL_16028739 (BA.5.11) by 11 nucleotide substitutions. The CM cluster consisted of two haplotypes, namely EPI_ISL_16093062 and EPI_ISL_16029195, with 349 and 857 isolates, respectively. The spike DNA sequence of EPI_ISL_16029195 differed from that of EPI_ISL_16028774 (BQ.1.1) by 13 nucleotide substitutions. The XBB cluster consisted of 14 haplotypes, with its diversifying center consisting of two haplotypes with GISAID accession IDs (EPI_ISL_) of 16044705 and 16206019. Among these haplotypes, the spike protein haplotype of EPI_ISL_16044705 was the most abundant, with 24144 isolates recorded. It differed from the haplotype of EPI_ISL_16040256 (BN.1.2) by 13 nucleotide substitutions and from the haplotype of EPI_ISL_16028739 (BA.5.11) by 22 substitutions. The DNA haplotype of EPI_ISL_16168343 (XBC.1) differed from that of EPI_ISL_15973011 (BA.5.2) by 19 nucleotide substitutions.

**Fig. 3.**
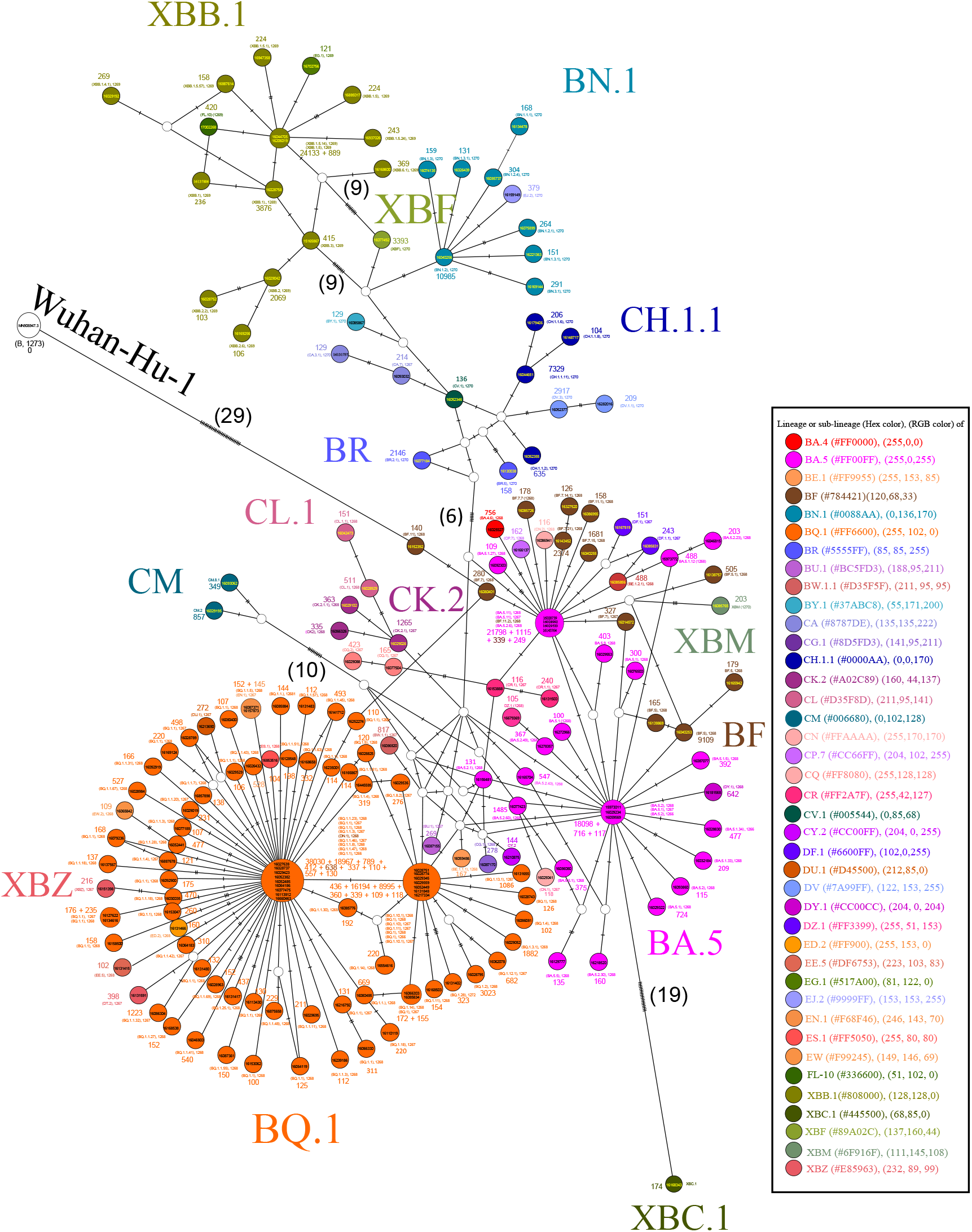
Median-join network of SARS-CoV-2 spike DNA sequences from December 2022 to February 2023. GISAID ID was labeled inside the circles. The number of isolates and lineages were labeled outside the circles. The number of nucleotide substitutions between haplotypes was labeled on the lines with hatch bars. When the hatch bars exceed 5, the substitutions were also labeled with numbers.

### Positive selection sites of spike protein

The values of likelihood ratio tests of M0 versus M3, M1a versus M2, and M7 versus M8 comparisons were larger the critical values at 0.01 level. The results suggest that the M3, M2, and M8 models were statistically better than M0, M1a, and M7 models, respectively. The Bayes empirical Bayes (BEB) analyses of M2 models identified the 25 amino-acid sites under positive selection. These sites were located at the positions of L5**, W152 **, G181 **, N185 **, G213*, V213*, H245*, Y248*, D253**, S255*, S256**, G257*, R346**, R408*, K444**, V445**, G446**, N450**, L452**, N460*, F486**, Q613**, Q675*, T883**, P1162**, and V1264**, which were statistically significant at 0.05 (*) and 0.01 (**) levels. The M8 model identified an additional one more site at V83* which was not identified by M2 model (Table 2). The site of L5 was located in signal peptide domain (SP) of the spike protein. The V83, W152, G181, N185, G213, H245, Y248, D253, S255, S256 and G257 were located in N-terminal domain (NTD). The R346, R408, K444, V445, G446, N450, L452, N460, and F486 were located in receptor binding domain (RBD). The Q613 and Q675 were located in C-terminal domain 2 (CTD2). The T883 was located in fusion-peptide proximal region (FPPR). The P1162 was located between HR1 and HR2. The V1264 was located in cytoplasmic tail (CT). The nonsynonymous substitutions of selection sites ranged from 4 to 11 in each protein haplotype and the same nonsynonymous substitution in the same selection sites usually occurred in different lineages except the substitutions of V83A, V213E, and V445P were exclusively occurred in the XBB lineage. Among these selection sites, the site of 444 had the largest amino acid diversity. The nonsynonymous substitutions included K444R, K444T, K444M, and K444N that were occurred in 7, 94, 4, 5 of 188 protein haplotypes, respectively.

**Table 1.** The net average genetic distances of per sequence between nine-lineage spike proteins. All ambiguous positions were removed for each sequence pair (pairwise deletion option).

|  | Wuhan-Hu-1 | B.1.1.529.2<br>(BA.2) | B.1.1.529.4<br>(BA.4) | B.1.1.529.5<br>(BA.5) | XBB | XBC | XBF | XBM |
| --- | --- | --- | --- | --- | --- | --- | --- | --- |
| B.1.1.529.2<br>(BA.2) | 34.54 |  |  |  |  |  |  |  |
| B.1.1.529.4<br>(BA.4) | 31.00 | 9.41 |  |  |  |  |  |  |
| B.1.1.529.5<br>(BA.5) | 37.07 | 7.30 | 1.99 |  |  |  |  |  |
| XBB | 36.62 | 11.87 | 13.18 | 11.73 |  |  |  |  |
| XBC | 35.00 | 16.71 | 15.00 | 14.12 | 19.62 |  |  |  |
| XBF | 37.00 | 1.71 | 12.00 | 10.52 | 11.62 | 18.00 |  |  |
| XBM | 33.00 | 11.41 | 4.00 | 3.90 | 15.18 | 17.00 | 14.00 |  |
| XBZ | 31.00 | 8.67 | 4.00 | 1.40 | 12.93 | 17.00 | 12.00 | 6.00 |

**Table 2.**
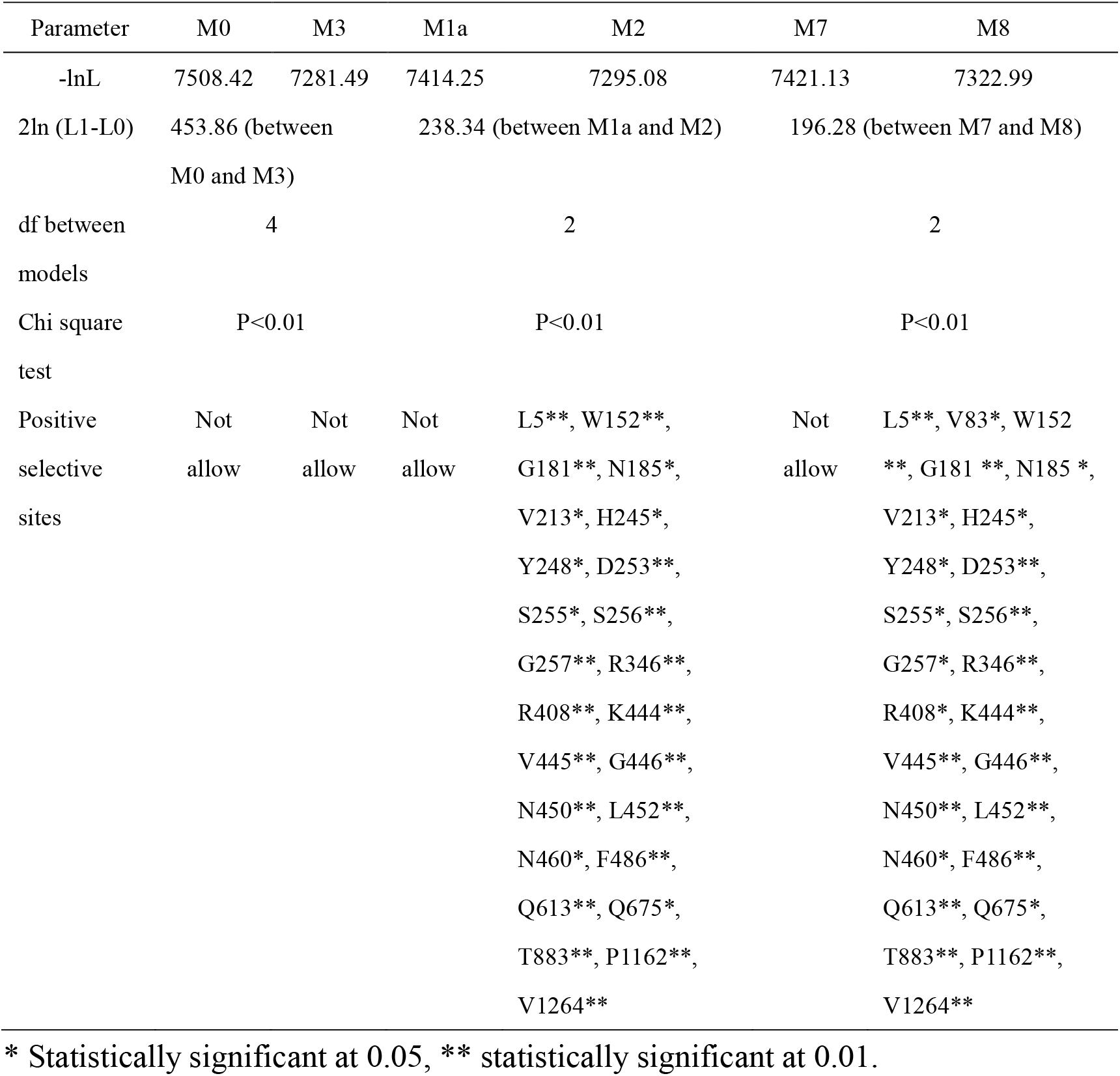
Likelihood ration test of M0 vs M3, M1a vs M2, M7 vs M8, and amino acid site of spike protein under positive selection.

| Parameter | M0 | M3 | M1a | M2 | M7 | M8 |
| --- | --- | --- | --- | --- | --- | --- |
| -lnL | 7508.42 | 7281.49 | 7414.25 | 7295.08 | 7421.13 | 7322.99 |
| 2ln (L1-L0) | 453.86 (between M0 and M3) |  | 238.34 (between M1a and M2) |  | 196.28 (between M7 and M8) |  |
| df between models | 4 |  | 2 |  | 2 |  |
| Chi square test | P<0.01 |  | P<0.01 |  | P<0.01 |  |
| Positive selective sites | Not allow | Not allow | Not allow | L5**, W152**, G181**, N185*, V213*, H245*, Y248*, D253**, S255*, S256**, G257**, R346**, R408**, K444**, V445**, G446**, N450**, L452**, N460*, F486**, Q613**, Q675*, T883**, P1162**, V1264** | Not allow | L5**, V83*, W152**, G181 **, N185 *, V213*, H245*, Y248*, D253**, S255*, S256**, G257*, R346**, R408*, K444**, V445**, G446**, N450**, L452**, N460*, F486**, Q613**, Q675*, T883**, P1162**, V1264** |

\* Statistically significant at 0.05, \*\* statistically significant at 0.01.

## Discussion

The presence of long and short spike protein sequences, with variations in amino acid length compared to the original Wuhan-Hu-1 spike protein, is not completely due to incomplete sequencing or sequence error. This conclusion is based on several observations made in the study. Firstly, the sequences did not contain ambiguous amino acids (represented by X). Secondly, all the sequences analyzed contained a start codon (M), Additionally, most sequences had a complete C-terminus domain. It is important to note that these variations in amino acid length were typically observed in the signal peptide or N-terminus domains, and rarely in the S2 region. Importantly, these insertions and deletions were never observed in the receptor binding domain (RBD) of the spike protein. The RBD is responsible for binding to the ACE-2 receptor, which is essential for viral entry into host cells. The fact that strains with long or short spike proteins still maintained infectivity suggests that they were still able to bind to the ACE-2 receptor despite these variations of sequence lengths.

The results showed that the net average genetic distances of spike protein between the Wuhan-Hu-1 strain and lineages of B.1.1.529.2, B.1.1.529.4, B.1.1.529.5, XBB, XBC, XBF, XBM, and XBZ ranged from 30.07 (between Wuhan-Hu-1 and BA.1.1.529.5) to 37 (between Wuhan-Hu-1 and XBF) amino acids per sequence. The results showed there was a great difference between the original (Wuhan-Hu-1) and current strains. Furthermore, the study specifically mentions the genetic distances between the XBB strain and several other strains. The genetic distances between XBB and lineages of B.1.1.529.2, B.1.1.529.4, B.1.1.529.5, XBC, XBF, XBM, and XBZ were 11.87, 13.18, 11.73, 19.62, 11.62, 15.18, and 12.93, respectively. Among these strains, XBC had the largest difference from XBB, with 19.62 amino acids per sequences. XBC was a recombinant of BA.2 Omicron (the most mutated) and B.1.617.2 Delta (the most severity) strains [22, 23]. It is important to continue surveillance and monitor the evolution of XBC.

The results of phylogenetic tree (Fig. 2) and median-join network (Fig. 3) revealed that the presence of multiple lineages of SARS-CoV-2 during December 2022 to February 2023. However, the majority of these lineages were descendants of three major lineages: BA.2, BA.5, and XBB. To help summarize the relationships between the lineages, the study employed the use of simplified names based on the Pango lineage nomenclature. However, the full names providing a more detailed and precise identification of the lineages.

Firstly, the BA.2 (BA.1.1.529.2) consisted of the sub-lineages of CM (B.1.1.529.2.3.20), CA (B.1.1.529.2.75.2), CV (B.1.1.529.2.75.3.1.1.3), DV (B.1.1.529.2.75.3.4.1.1.1.1.1), CH (B.1.1.529.2.75.3.4.1.1), BR (B.1.1.529.2.75.4), BN (B.1.1.529.2.75.5), EJ.2 (B.1.1.529.2.75.5.1.3.8.2) and BY (B.1.1.529.2.75.6).

Secondly, the BA.5 (BA. 1.1.529.5) consisted of eight major sub-lineages, i.e., BA.5.1, BA. 5.2, BA.5.3, BA.5.5, BA.5.6, BA.5.9, BA.5.10, and BA.5.11 in this study. The descendants of BA.5.1 (B.1.1.529.5.1) consisted of BA.5.1.5 (B.1.1.529.5.1.5), BA.5.1.12 (B.1.1.529.5.1.12), BA.5.1.27 (B.1.1.529.5.1.27), and CL.1 (B.1.1.529.5.1.29.1) in this study. The descendants of BA.5.2 consisted of BA.5.2.1 (B.1.1.529.5.2.1), BF.5 (B.1.1.529.5.2.1.5), BF.7 (B.1.1.529.5.2.1.7), BU.1 (B.1.1.529.5.2.16.1), CR.1.1 (B.1.1.529.5.2.18.1.1), CR.1.2 (B.1.1.529.5.2.18.1.2), CN.1 (B.1.1.529.5.2.21.1), CN.2 (B.1.1.529.5.2.21.2), BA.5.2.23 (B.1.1.529.5.2.23), CK.2 (B.1.1.529.5.2.24), in this study. The descendants of BA.5.3 (B.1.1.529.5.3) consisted of BQ.1 (B.1.1.529.5.3.1.1.1.1.1), DU.1 (B.1.1.529.5.3.1.1.1.1.1.1.2.1), and CQ (B.1.1.529.5.3.1.4.1.1) in this study. The descendants of BA.5.6 (B.1.1.529.5.6.) consisted of BW.1.1 (B.1.1.529.5.6.2.1.1) in this study. The descendants of BA.5.10 ((B.1.1.529.5.10) consisted of DF (B.1.1.529.5.10.1) in this study. The BA.5.11 consisted of the BA.5.11 only. Thirdly, XBB was the recombinant of two BA.2 lineages, i.e., BJ.1 and BM1.1.1 [24]. The EG.1 and FL.10 were the abbreviations of XBB.1.9.2.1 and XBB.1.9.1.10, respectively. The other recombinants include XBC (a recombinant of BA.2 Omicron and Delta), XBF (a recombinant of BA.5 and BA.2.75), and XBZ (a recombinant of BA.5.2 and EF.1.3) based on the Covid-lineage Pango designation (Roemer, 2022) [23]. The most dominant variant was the strain BQ. 1.1.23 with the representative accession number of EPI_ISL_16027638, and had 55919 identical isolate sequences, following by XBB.1.5 (representative accession number EPI_ISL_16044705, 24133 identical isolate sequences), and BA.5.11 (representative accession number EPI_ISL_16028739, 21798 identical isolate sequences) during December, 2022 to February, 2023.

The previous study demonstrated that certain mutations in the receptor-binding domain (RBD) of the spike protein, specifically at positions R346, K356, K444, V445, G446, N450, L452, N460, F486, F490, R493, or S494, could lead to the evasion of neutralizing monoclonal antibodies (mAbs) or enhance binding to the ACE2 receptor (Cao et al., 2022). In the present study, we found that mutations at R346, K444, V445, G446, N450, L452, N460, and F486 had a nonsynonymous versus synonymous substitution ratio greater than 1, indicating positive selection. This suggests that these sites were undergoing evolutionary changes that may confer selective advantages to the virus. However, mutations at K356, F490, F493, or S494 did not exhibit a nonsynonymous versus synonymous substitution ratio greater than 1 in the present analysis, suggesting that these sites were not under positive selection during the specific time frame examined (December 2022 to February 2023) in the study (Table 2). This finding contrasts with the previous study, which analyzed sequences from January 2021 to October 2022. I propose that the discrepancy in results between the previous and present studies may be attributed to antigenic shift. It’s possible that the evolutionary dynamics and selective pressures acting on SARS-CoV-2 may have shifted, leading to different mutations being favored in different time periods. Additionally, the present study identified positive selection for mutations occurring outside of the RBD domain. These sites included L5, V83, W152, G181, N185, G213, H245, Y248, D253, S255, S256, Q613, Q675, T883, P1162, and V1264. However, the effects of these mutations on the fitness of SARS-CoV-2 remain to be investigated.

In this study, it was observed that multiple strains coexisted between December 2022 and February 2023. However, the majority of these strains belonged to the lineages or sub-lineages of BA.2 (BA.1.1.529.2), BA.5 (BA.1.1.529.5), and XBB (Fig. 2). The diversifying centers of BN.1.2, BQ.1, BA.5.11, XBB were the isolate sequences with representative accession IDs (EPI_ISL_) of 16040256, 16027638, 16028739, and16044705, respectively (Fig. 3). I propose that the complete sequences or the receptor binding domain of these spike DNA sequences could be potential candidates for vaccine design. This suggests that these sequences may possess important characteristics that can be utilized in the development of effective vaccines against SARS-CoV-2.

As of June 10, 2023, just before submitting our manuscript, the XBB.1.5 and XBB.1.16 strains have emerged as the globally dominant strains, with respective frequencies of 72% and 12% based on data from GISAID [25]. These strains have gained prominence and become widespread within the population. Additionally, the XBC variant is a recombinant of BA.2 (Omicron) and B.1.617.2 (Delta) [17, 23].

XBC exhibits significant differences from the XBB lineages and its sub-lineages, making it a distinct variant from XBB. Considering the success of the Omicron variant [3, 4], I propose that the XBC.1 strain or its sub-lineages could potentially become dominant strains following the XBB.1 lineage and its sub-lineages. Continued surveillance and research are necessary to monitor the evolution and potential impact of these variants on public health.

## Acknowledgements

We gratefully acknowledge all data contributors, i.e., the Authors and their Originating laboratories responsible for obtaining the specimens, and their Submitting laboratories for generating the genetic sequence and metadata and sharing via the GISAID Initiative, on which this research is based (Supplementary Table 1).

## Availability of data and materials

All sequences were downloaded from Global Initiative on Sharing Avian Influenza Data (GISAID, https://www.gisaid.org/) and GenBank (https://www.ncbi.nlm.nih.gov/nucleotide/).

## Ethics approval and consent to participate

Not applicable.

## Consent for publication

Not applicable.

## Author details

^1^Department of Life Sciences, National Chung Hsing University, Taiwan, R.O.C. 145 Xingda Road., South District., Taichung City 40227, Fax: +886-4-22874740 Tel: Tel: +886-4-22840416 *Correspondence: Hsiao-Wei Kao

